# Bending of the BST-2 coiled-coil during viral budding

**DOI:** 10.1101/160242

**Authors:** Kadir A. Ozcan, Christopher E. Berndsen

## Abstract

BST-2/tetherin is a human extracellular transmembrane protein that serves as a host defense factor against HIV-1 and other viruses by inhibiting viral spreading. Structurally, BST-2 is a homodimeric coiled-coil that is connected to the host cell membrane by N and C terminal transmembrane anchors. The C-terminal membrane anchor of BST-2 is inserted into the budding virus while the N-terminal membrane anchor remains in the host cell membrane creating a viral tether. The structural mechanism of viral budding and tethering as mediated by BST-2 is not clear. To more fully describe the mechanism of viral tethering, we created a model of BST-2 embedded in a membrane and used steered molecular dynamics to simulate the transition from the host cell membrane associated BST-2 and the cell-virus membrane bridging form. We observed that BST-2 did not transition as a rigid structure, but instead bent at sites with a reduced interface between the helices of the coiled-coil. The simulations for the human BST-2 were then compared with simulations on the mouse homolog, which has a more stable coiled-coil. We observed that the mouse homolog spread the bending across the ectodomain, rather than breaking at discrete points as observed with the human homolog. These simulations support previous biochemical and cellular work suggesting some flexibility in the coiled-coil is necessary for viral tethering, while also highlighting how subtle changes in protein sequence can influence the dynamics and stability of proteins with overall similar structure.

## Introduction

BST-2 (also known as CD317 or Tetherin) is a human cell membrane embedded protein which is associated primarily with tethering a diverse group of viruses to the host cell membrane^1^. While BST-2 also functions in organizing the cytoskeleton and signaling, the antiviral activities are best characterized ^1–3^. Structurally, BST-2 has four main features: a cytoplasmic region, an N-terminal transmembrane helix, an extracellular coiled coil domain, and a C-terminal membrane anchor which is either a GPI anchor or a protein based anchor depending on the species^4–8^. BST-2 is a constitutive dimer and the extracellular coiled coil intertwines the two monomers connected by at least one disulfide^9,10^. Finally, the protein is known to be glycosylated on at least one of two asparagine residues within the ectodomain which is important for trafficking the protein to the cell membrane^10^.

Several cellular and biochemical studies have described aspects of the mechanism of viral tethering by BST-2^10–14^. Andrew and coworkers showed that the size of the ectodomain was not strictly defined and that one of the three disulfides must be intact for effective viral tethering^10,15^. Welbourn and coworkers showed that the disulfides could be moved to several positions within the ectodomain if the structure of the coiled-coil was not disrupted^9^. These disulfides appear to stabilize the structure of the coiled-coil and make it more resistant to unwinding^9,13^. Thus, the disulfides show a critical role in viral tethering by increasing the strength of the tether. Mechanistically, several proposals were described for how BST-2 operated to hold the virus in place until Venaktesh and Bienasz showed that the C-terminal anchor attached to the budding virus forming a bridge between the two membranes^11^. This mechanism was later supported by Strauss and coworkers who characterized the 3-D structure of tethered viruses and the host cells via electron microscopy^16^. Measurements of the gap between the virus and the host cell match the length of the ectodomain of BST-2 supporting a mechanism where a single BST-2 dimer forms a linear bridge between the cell and the viral particle^11,16^.

The description of how BST-2 tethers viruses has not however been extended to show how BST-2 transitions from the cell membrane bound to the bridging position. This transition would occur as the virus is budding but not fully detached from the membrane and would be influenced by the structure of the ectodomain. Here, we study this aspect of BST-2 function using molecular modeling and dynamics of the human and murine BST-2 homologs. We observed the structural changes in the ectodomain of BST-2 as it was pulled away from the membrane finding sequence encoded instabilities allow the ectodomain to bend during this transition. The murine BST-2 homolog showed similar structural transitions to the human homolog but the bend sites were distinct from the human homolog as expected given the low sequence conservation. These findings provide an atomic level description of viral tethering for two BST-2 homologs suggesting despite the absence of sequence conservation a conserved set of structural transitions to the bridging form of the protein.

## Materials and Methods

### Modeling of human and murine BST-2 in a lipid membrane

The structure of the ectodomain of human BST-2 (PDB entry 3MQB) and the structure of the BST-2 N-terminal anchor were used to create the model of full length human BST-2 in YASARA 16.9.23 ^17,18,6,19^. The N-terminal transmembrane domain spanned from residues 24-46 and was connected to the ectodomain by a coil with residues 47-50. The C-terminal transmembrane domains were created in YASARA 16.9.23 as alpha helices by using the sequence from Uniprot (Q10589). The C-terminal transmembrane domain spanned from residues 168-180 and was connected to the C-terminal of the ectodomain by a coil containing residues 154-168. All simulations were equilibrated in an explicit solvent at 298 K, 0.9% NaCl, and pH 7.4, using the AMBER14^20^. The full-length BST-2 with transmembrane domains was equilibrated into a phospholipid membrane consisting of 100% phosphotidyethanolamine (PEA) using the Build Membrane macro in YASARA. The per residues RMSF values were calculating using the internal trajectory tools within YASARA.

The structure of the murine ectodomain was from PDB entry 3NI0 and N-terminal and C-terminal loops and membrane anchors built in YASARA using the sequence of murine BST-2 found in Uniprot entry Q8R2Q8^8^. The model was equilibrated using the procedure described for the human BST-2 model. Membrane anchoring regions were approximated using the similarity between the human and mouse sequences.

### Steered Molecular Dynamics simulations

Steered molecular dynamics (SMD) was performed using the previously created full length BST-2 in membrane with the ectodomain cleaved at Ser 163 to allow for the ectodomain of BST-2 to be pulled away from the membrane. A force of 25 pN was applied to Ser162. Simulations conditions were 298 K, pH 7.0, and 0.9% NaCl in explicit solvent in AMBER14. Snapshots were recorded every 0.01 ns until the ectodomain was separated fully from the lipid membrane. The psi angles in each simulation snapshots were measured using a self-created macro and data were compiled into a .csv file using another self-created macro. Data were plotted using Excel and Plotly. Heat maps were generated on the psi angles based on degree of alpha helicity. In SMD experiments on the murine BST-2 homolog, the ectodomain was cleaved at amino acid Ser 154 and experiments then proceeded as described for the human homolog.

## Results

The structure of BST-2 contains a N-terminal transmembrane anchor, attached to an extracellular coiled-coil domain, and a C-terminal anchor. Up to this point, the structure of full-length BST-2 has not yet been fully described. Only the structure of the N-terminal transmembrane anchor and the ectodomain have been determined experimentally^6,8,19,21,22^. We modeled the full-length structure of BST-2 embedded in a lipid membrane to detail the mechanism of viral tethering at an atomic level. To build the model of the human protein, we used the existing structure of the ectodomain and N-terminal transmembrane anchor, which we connected by a flexible loop^6,19^. There is evidence that the C-terminal anchor of human BST-2 is protein rather than the GPI anchor observed in the rat homolog of BST-2^23^. Therefore, we modeled the C-terminal anchor *ab initio* as an alpha helix and connected it to the ectodomain via a flexible loop consisting of residues Tyr154 to Ile168 (Figure 1A). Once the model of BST-2 was built, we then added a membrane around the protein to determine the effects of anchoring the protein in a lipid membrane. The membrane was composed of phosphatidylethanolamine (PEA) and the anchors were manually placed in to the membrane followed by equilibration in the membrane. After 50 ns of equilibration, the ectodomain and membrane anchors within the model maintained a helical structure suggesting the overall model was chemically sound (Figure 1A and B).

**Figure 1.**
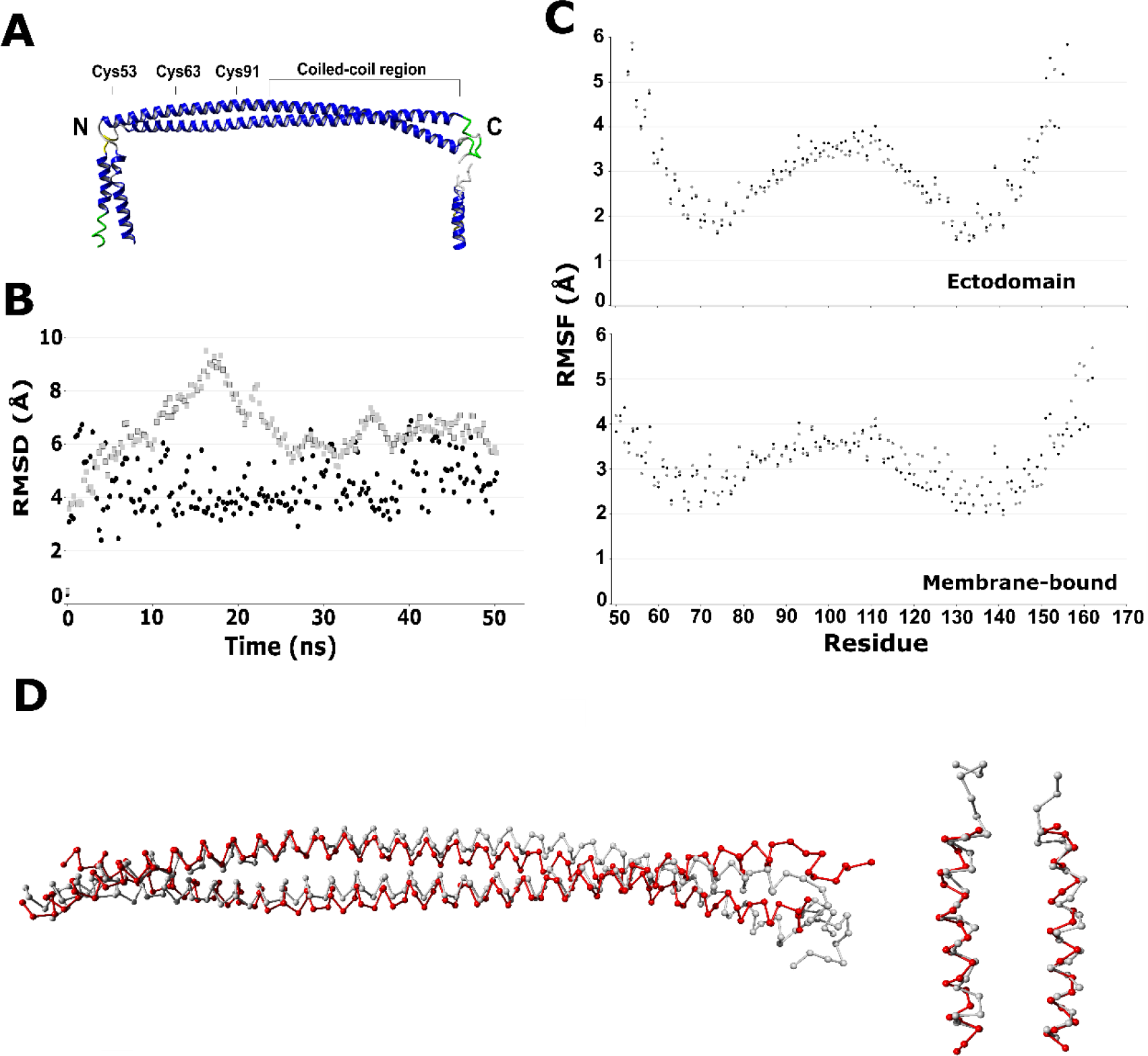
Molecular dynamics of BST-2 embedded in the membrane. (A) Model of full-length BST-2 (B) Plot of RMSD vs. time for membrane bound, full-length BST-2 (grey squares) and the ectodomain alone from structure 3MQB (black circles) (C) Plot of RMSF vs. residue number for membrane bound, full-length BST-2 (bottom panel) and the ectodomain alone from structure 3MQB (top panel). Each molecule of the dimer is shown with a different color. (D) Alignment of the 3MQB crystal structure of the BST-2 ectodomain of the free (red) and average structure of the ectodomain from the membrane bound protein (white) after equilibration. The RMSD value was 5.8 Å. Right panel shows the 2LK9 NMR structure of the BST-2 N-terminal domains (red) compared to the average structure of the N-terminal domains from the membrane bound protein (white) after equilibration. The RMSD values are 1.4 and 1.8 Å

Because a structure or model of BST-2 embedded in the lipid membrane has not previously been described, we observed the effect of the membrane on the ectodomain dynamics and compared these data to equilibration of the ectodomain in the absence of a lipid membrane (Figure 1B-E). For our comparison, we calculated the Root Mean Square Fluctuation (RMSF) for each residue in the model, which reflects the dynamics of the amino acid. For residues in the ectodomain of the membrane bound model, a plot of RMSF vs. residue number showed a “W” shape (Figure 1C). The maximum value excluding the termini was centered around amino acid 105 and the two minima were located around amino acids 75 and 135. The most dynamic areas were the loops at the end of the ectodomain and for residues 95-115 which are in the beginning of the canonical coiled-coiled structure^15^. Molecular dynamics simulations of the ectodomain crystal structure 3MQB show a similar “W” shape that aligns well with the membrane bound model (Figure 1C). Alignment of the average membrane bound structures with the crystal structure of the ectodomain or the NMR structure of the N-terminal anchor show the equilibrated structure aligns well with experimentally determined structures and the primary differences are observed at the C-terminus (Figure 1D). These data suggest that the membrane does not alter the overall structure or dynamics of the BST-2.

As BST-2 is an extracellular transmembrane anchored protein, the protein is glycosylated upon exit from the ER^10^. The structural implications of glycosylation of BST-2 are an additional factor to consider as there is a possibility that the complex glycosylation may alter the structure and dynamics of the ectodomain. To understand structural implications of glycosylation, Asn65 and Asn92 of the membrane bound model of BST-2 were glycosylated with complex glycosylation as seen in Supplemental Figure 1A. This glycosylation added a mass of about 4.4kDa to both monomers of the ectodomain dimer. Equilibration of the glycosylated BST-2 in membrane yielded an RMSF plot showing the same shape and levels of dynamics as the non-glycosylated membrane bound and ectodomain structures of BST-2 (Supplemental Figure 1B and 1C). The structures of the modified and unmodified ectodomain aligned well suggesting no structural changes (Supplemental Figure 1D). The similar RMSD and RMSF values are indicative of similar structure and dynamics of the ectodomain. These data support the idea that the intrinsic dynamics of BST-2 are not influenced by post-translational modification. Moreover, the structure of the protein in the membrane

Previous cellular work has shown that during viral tethering BST-2 bridges the gap between the host cell and the viral particle^11,16^. The distance spanned by BST-2 in microscopy studies is consistent with the length of the ectodomain, suggesting a simple transition of the ectodomain from the membrane bound form to the bridging form as shown in Figure 2A. However, the structural mechanism for moving between the two states is not clear. Therefore, we simulated the transition from the membrane bound to the host cell-viral particle bridging forms. In a previous study, we showed that a steered molecular dynamics simulation pulling the termini of the BST-2 ectodomain to mimic viral tethering correlated well with biochemical experiments^13^. Therefore, we turned to steered molecular dynamics to simulate the conformational changes BST-2 during the transition to the bridging form using our model of the membrane bound, full-length BST-2. For the simulation, we cleaved the loop between the ectodomain and the C-terminal membrane anchor at Ser163. We then applied 25 pN of force to Ser162 of both molecules of the BST-2 dimer such that the C-terminal of the ectodomain was pulled away from the membrane. The simulation setup is illustrated in Figure 2A. The steered simulations were run until the ectodomain was completely separated from the lipid membrane and nearly perpendicular to the membrane (Figure 2B). We used this approach to reduce the complexity of the simulation to a computationally manageable scenario that did not include simulating viral protrusion and lipid membrane cleavage. The final orientation of the ectodomain would put the C-terminal anchor in the virus and the N-terminal anchor in the host cell, which is consistent with cellular studies^11^.

**Figure 2.**
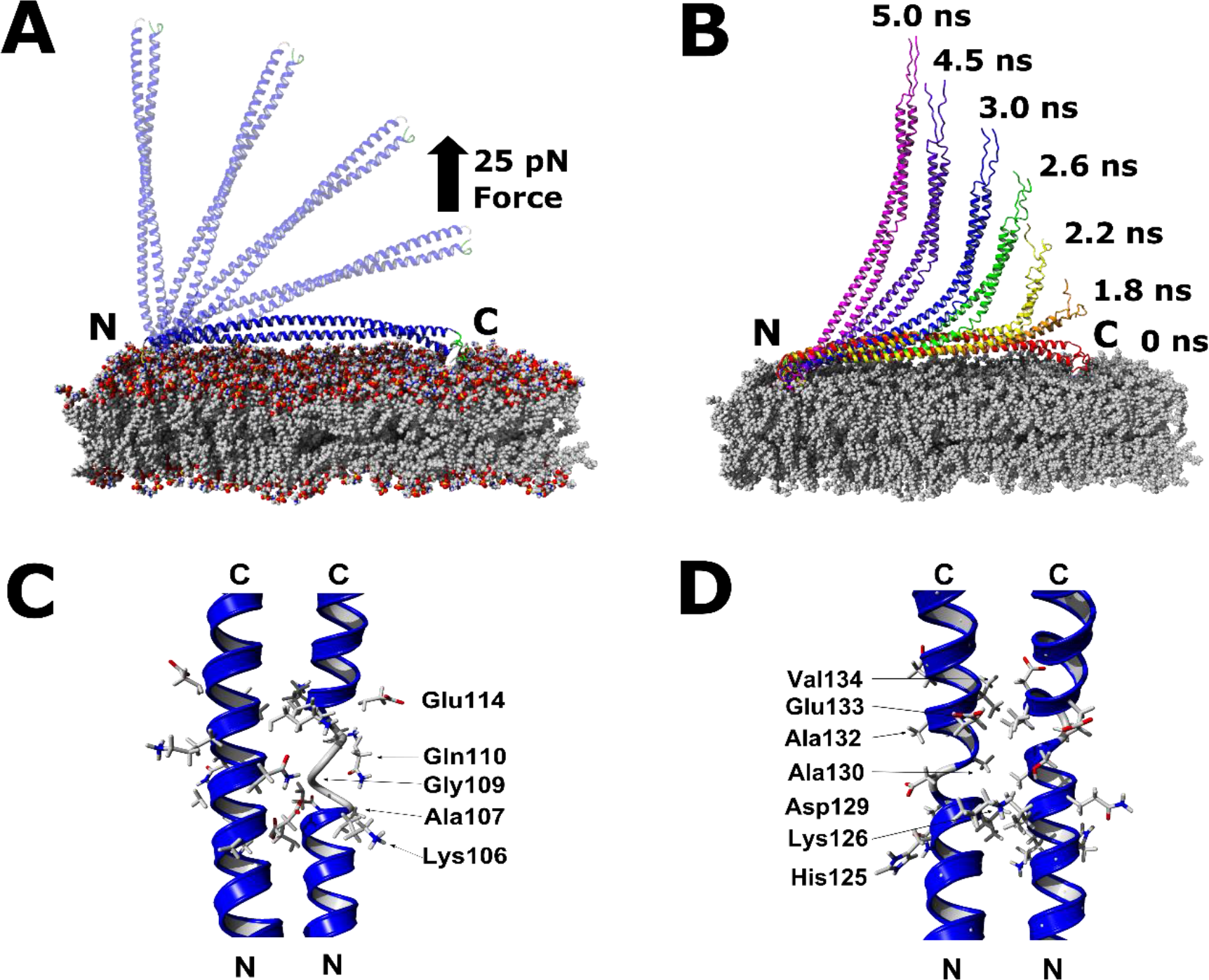
Steered molecular dynamics on BST-2 transitioning from the membrane bound to bridging position. (A) Experimental setup for SMD on BST-2 (B) Selected structures from SMD experiments (C) Structure showing reduced dimer interface around Gly109 (D) Structure showing reduced dimer interface around Ala130

Initially, we hypothesized that the BST-2 ectodomain would move away from the membrane rigidly as a single coiled-coil unit with minimal bending of the structure. This movement can be thought of as a door (BST-2) swinging away from the membrane (Figure 2A). For the first 1.5 ns of the simulation, the ectodomain remained near the surface of the membrane. The only changes were in the C-terminal loop, as this flexible region pulled away from the membrane. After 1.5 ns, the ectodomain began to move away from the membrane at a linear rate. By 3.7 ns, BST-2 was oriented nearly perpendicular to the membrane and the simulation was stopped (Figure 2B). A second simulation with the same set up performed to confirm reproducibility of the simulation (Supplemental Figure 2). In both simulations, BST-2 moved not as a single rigid unit but the coiled-coil domain bent ∼45° relative to the starting structure (Figure 2B). The clear breakpoints in the coiled-coil were localized around Gly109 and Ala130 (Figure 2C and 2D). The ectodomain peeled away from the membrane in three segments: 1) Ser50 to Gly109, 2) Gly109 to Ala130, and 3) Ala130 to Ser162. Residues 130 to 162 lifted first, followed by residues 109 to 130 and finally the N-terminal region consisting of residues 50 to 109 (Figure 2B). We next analyzed the psi angles of BST-2 to observe the duration and extent of the bonding changes during the simulation (Figure 3A). The psi angle analysis shows that the alpha helical structure is maintained during the simulation except for localized disruptions of the helix around residues Ala130 and Gly109 (Figure 3A and Supplemental Figure 2A). These break points in the coiled-coil were consistent in both simulations. Analysis of the bending angle of the coiled-coil and the three segments shows that as the distance between the membrane and C-terminal rapidly increases after 1.5 ns, that bends in the coil occur at Gly109 and Ala130 while the ectodomain between residues 50 and 109 maintains a consistent angle (Figure 3C and Supplemental Figure 2B). By the end of the simulation, the angles are returning to values near to the beginning of the simulation, indicating re-formation of the alpha helix (Figure 3C and Supplemental Figures 2B and 2C).

**Figure 3.**
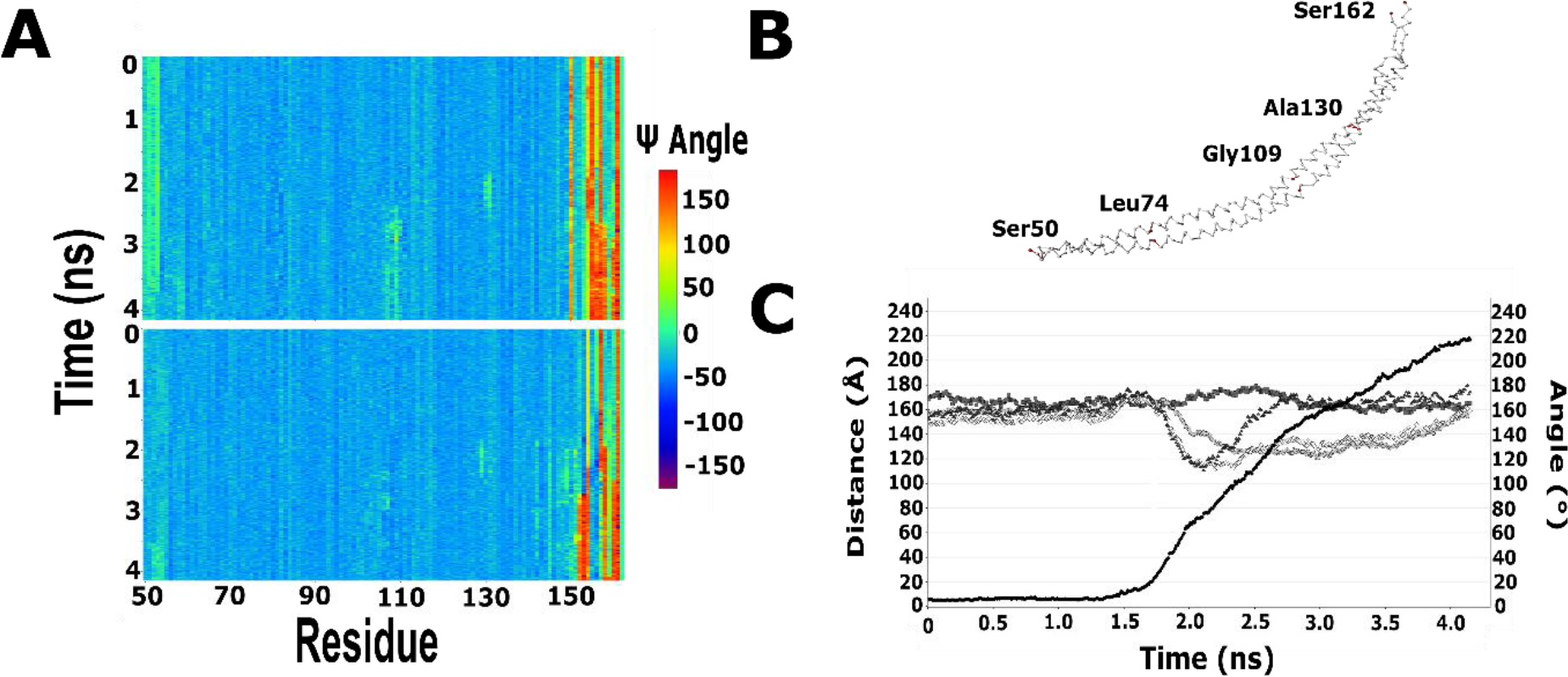
Structure analysis of the steered molecular dynamics experiment. (A) Heatmap showing the psi angle (Cα-CO) for each amino acid at each 0.01 ns time frame. Alpha helices have psi angles of approximately −60° (B) Labeled diagram of BST-2 ectodomain (C) Plot of distance or angle versus time for the steered molecular dynamics experiment of BST-2. Distance is shown in solid black circles, the angle between amino acids 50, 74, and 109 is shown in grey squares, the angle between amino acids 50, 109, and 162 are shown in grey circles, the angle between 50, 130 and 162 is shown in white diamonds, and the angle between 109, 130, and 162 is shown in grey triangles.

The breakdown of the alpha helical structure at Gly109 and Ala130 in the BST- 2 ectodomain led us to search for the molecular basis for the weakness (Figure 3A). In a coiled-coil, the interface residues are typically leucine, isoleucine or valine, thus deviations from these residues can affect coil stability^24^. Analysis of the dimer interface in the human BST-2 crystal structure shows a hole in between the monomers at Gly109 suggesting reduced interaction between the two monomers around this site (Figure 2C). Ala130 is also at the interface between the two helices and the smaller side chain of alanine could affect the interaction (Figure 2D). Therefore, bending of the coiled-coil seems dependent on the sequence amino acids at the interface of the monomers. For comparison, the only other BST-2 homolog with a structure is the mouse homolog so we aligned the crystals structures of the human and murine homologs (data not shown)^8,6^. We did not observe any gaps in the murine coil and note that melting temperature for the ectodomain of the murine BST-2 is higher than the value for the human homolog^8,13^. These data suggest that the mouse homolog has a more rigid dimer interface which might alter the structure of the ectodomain during the transition from membrane to bridging position.

We wanted to further explore the conservation of this mechanism and the link between dimer interface sequence and ectodomain bending in our steered simulations. To test whether the murine homolog shows a distinct set of bends and breaks within the coiled coil, we next modeled the structure of the membrane bound BST-2 from mouse and performed molecular dynamics experiments to those that we performed on the human protein. Murine BST-2 shows a W shaped RMSF vs. residue number plot after equilibration of the model within a lipid membrane however the shape is asymmetric with distinct local minima at Cys68 and Val 128 (Figures 4A-C). In SMD experiments, the murine protein bent 50 degrees relative to the starting angle but did not show a durable breaking of the helix (Figure 4D and 4E). Transient increases in the psi angles of the helix were observed at amino acids Ser 85, Ser 95, Asn 126, and Glu 134 over a similar time interval when the angle of ectodomain bending peaks (Figure 4E and 4F). The murine BST-2 appeared to bend as one unit at several locations simultaneously before returning to a linear form at the end of the simulation (Figure 4F). The tighter coiled-coil interface of the murine BST-2 appears to prevent breakdown of the ectodomain helices and spread the deformations across the entire domain.

**Figure 4.**
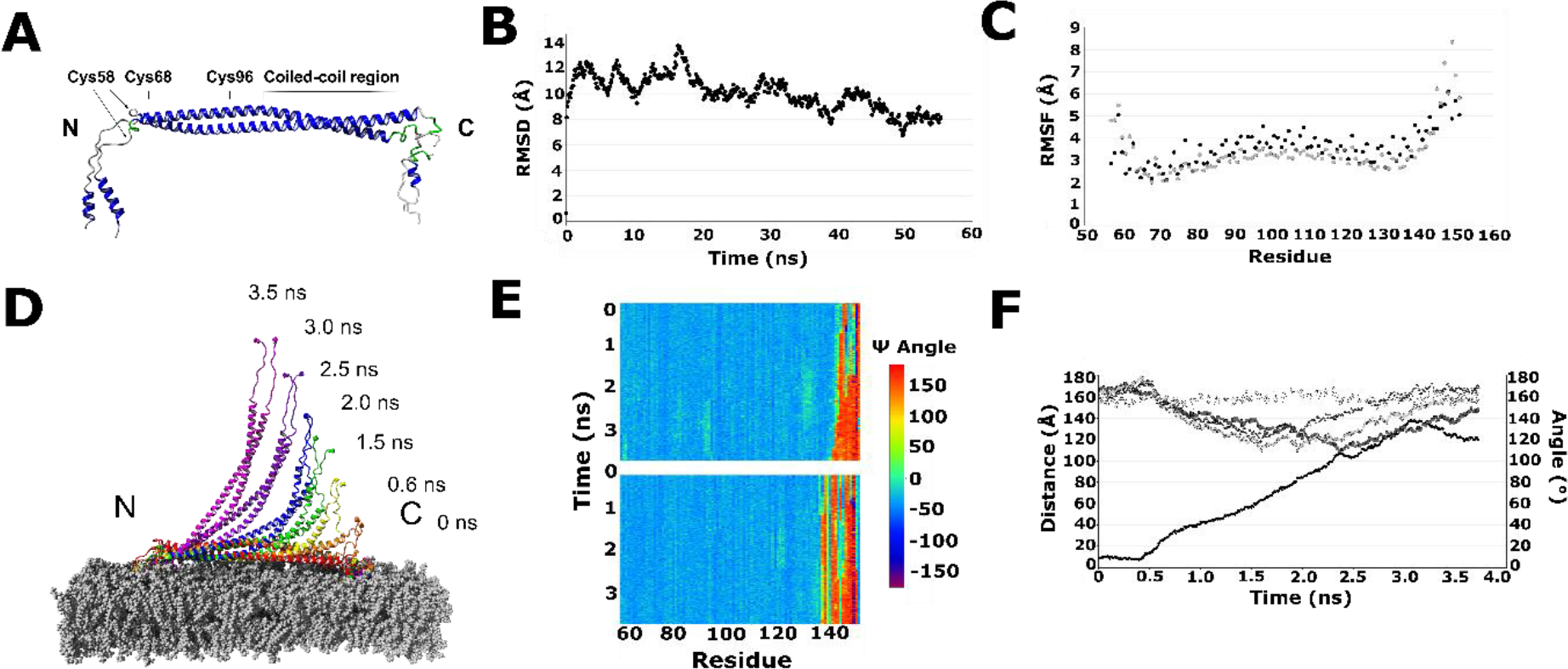
Model and steered molecular dynamics on murine BST-2. (A) Model of full-length murine BST-2 (B) Plot of RMSD vs. time (C) Plot of RMSF vs. amino acid number (D) Selected structures from SMD experiments (E) Heatmap showing the psi angle (Cα-CO) for each amino acid at each 0.01 ns time frame. Alpha helices have psi angles of approximately −60° (F) Plot of distance or angle versus time for the steered molecular dynamics experiment of murine BST-2. Distance is shown in solid black circles, the angle between amino acids 63, 84, and 94 is shown in open circles, the angle between amino acids 63, 128, and 154 are shown in grey circles, the angle between 63,84, 154 is shown in grey squares, and the angle between 94, 128, and 154 is shown in grey triangles.

## Discussion

The antiviral protein BST-2 physically tethers budding viruses to the host cell membrane preventing the particle from spreading and infecting other cells^25^. While the cellular details of this process have been long studied and described, the mechanistic details regarding viral tethering are only now coming to light^11,13,16^. Using steered molecular dynamics, we simulated the transition of human and murine BST-2 from the host cell bound form to the host-virus bridging form (Figures 2, 3, and 4). Encoded weaknesses in the human protein sequence, which reduce the interaction surface between the monomers, delineated segments of human BST-2 in the RMSF data (Figures 1C, 2C and 2D). These break points permit bending of the protein without global unfolding of the coiled-coil. The murine homolog lacks apparent weaknesses such as those observed in the human structures. Comparison of these data between the human and murine homologs, show differences in the RMSF plot suggesting less of a segmented motion to the coiled-coil, which we attribute to the increased interaction surface between the monomers (Figures 1C and 4C). Further comparison of the psi angles and bending angles across the simulation time, show a segmented pattern for the human homolog while, the murine homolog bends as a single unit with increases in the psi angle at several spots simultaneously along the coiled-coil (Figures 3A, 3C, 4E and 4F). Thus, the interface sequence of the coiled-coil appears to influence the lateral stability and bending of the protein.

The biological significance of the bending and weaknesses is not immediately clear. Using cellular studies on BST-2 mutants and truncations, Andrew and coworkers showed that substitution of the coiled-coil sequence of vimentin in addition to or in place of the BST-2 coiled-coil only was functional when the second coiled-coil region of vimentin was used^15^. This section of vimentin contains a “stutter” in the alpha helix sequence which may permit the bending like we observe in our simulations ^26,27^. A canonical coiled-coil from vimentin or a non-coiled coil substitution showed reduced or absent function in viral tethering assays^15^. Further work using cysteine substitutions within the coiled-coil region (96 to 150) including substitutions at Gly109 and Ala130, which would strengthen the interaction between monomers, showed reduced or no viral tethering activity and reduced levels of dimerization^9^. In contrast, Cys substitutions in the N-terminal portion of the ectodomain (residues 50 to 95), which in our simulations was moved as a single unbroken region, showed mixed results with some substitutions performing viral tethering similar to the wild-type and all were able to dimerize^9^. These data suggest that there is a functional importance to having a flexibility encoded in the coiled-coil.

For viral tethering, BST-2 flexibility appears to run counter to the function of stably tethering viruses to the cell membrane. Using sequence analysis, Swiecki and coworkers proposed encoded instabilities in the coiled-coil^8^. However, the unstable regions proposed by their study extended in the region between amino acids 50 and 100, which we find to be rigid in our current study (Figure 2C). In our previous work however, we showed that this region was susceptible to unfolding when the ends of the ectodomain were pulled in opposite directions, which was supported by spectroscopic and mutational studies^9,13^. The coiled-coil region from amino acids ∼101 to 155 unfolded only after the N-terminal region had completely unfolded^13^. In this study, unwinding within the coiled-coil is observed around Gly109 after the N-terminus had unfolded and the psi angle of Ala130 was consistently higher than neighboring amino acids. Our current data showing breakdown of the human ectodomain at these spots support these previous findings despite a number of experimental differences^13^.

## Conclusion

We conclude that the lateral strength and tensile strength of the ectodomain are divided properties in the human protein. The N-terminal region consisting of amino acids 50 to 100 appears to mediate tensile strength while C-terminal region consisting of amino acids 101 to 155 appears to mediate bending during viral tethering. While we noted differences in the structures of the human and murine homologs during pulling away from the membrane, it is known that the human protein can restrict viral spreading from certain cell types in mouse models^28^. However, human BST-2 cannot be expressed throughout the mouse without causing defects in growth^28^. Thus, the differences in the bending of the ectodomain do not appear to be significant to viral tethering in certain situations, however for other functions the role of BST-2 structure and dynamics is not known.

## Acknowledgements

We gratefully acknowledge helpful discussions with Kelly E. Du Pont and Dr. Isaiah Sumner on the technical aspects and manuscript. We acknowledge support from the National Science Foundation Research Experience for Undergraduates Grant (CHE-1461175) and the Department of Chemistry and Biochemistry at James Madison University.

